# The cytoplasmic scaffolding protein PCNA regulates NLRP3 inflammasome activation in macrophages

**DOI:** 10.1101/2025.07.08.663719

**Authors:** Chloé Lopes, Karen Aymonnier, Nicolas Cagnard, Géraldine Falgarone, Léa Tourneur

## Abstract

Key components of innate immune responses, the NLRP3 sensor, ASC adaptor and Caspase-1 effector form the NLRP3 inflammasome, whose assembly leads to both inflammatory cytokines secretion and cell death. Its priming and activation are regulated by numerous proteins including the serine/threonine kinase NEK7. PCNA is a nuclear scaffolding protein essential for DNA replication. In neutrophils, which are differentiated cells deprived of proliferative capacity, PCNA is exclusively cytoplasmic and regulates neutrophil functions through interactions with different protein partners. Among them, pro-Caspases-3/8/9/10 interact with PCNA to regulate apoptosis. We therefore hypothesize that cytoplasmic PCNA could serve as a regulatory platform for the NLRP3 inflammasome via its interaction with its components. In PMA-differentiated THP-1 macrophages, PCNA relocalized mainly in the cytoplasm. Furthermore, PCNA scaffold inhibitors inhibited IL-1β secretion, pro-Caspase-1 cleavage and pyroptosis-activating gasdermin D cleavage, induced by either nigericin, monosodium urate crystals or *Escherichia coli* bacteria. Nigericin-stimulation resulted in cytoplasmic PCNA binding to NEK7 and potentially NLRP3, and the p21 peptide, the highest competitive inhibitor of PCNA partner binding, disrupted PCNA-NEK7 interaction. Finally, use of bone marrow-derived macrophages from p21 knockout mice suggested that the cyclin-dependent kinase inhibitor p21 might be a natural repressor of the NLRP3 inflammasome. This work provides a proof-of-principle for the potential of disrupting PCNA interactions to regulate both canonical and non-canonical NLRP3 inflammasomes.

## 1 INTRODUCTION

Inflammasomes are multiprotein complexes, predominantly expressed in innate immune cells, activated by stress or danger signals (1). They contain a pattern recognition receptor (PRR) that senses danger/damage-associated molecular patterns (DAMPs) or pathogen-associated molecular patterns (PAMPs). Upon recognizing DAMPs/PAMPs, the sensor PRR assembles with the adaptor protein ASC (apoptosis-associated speck-like protein containing a CARD domain) leading to the recruitment and auto-activation of inflammatory effector pro-Caspase-1. Activated Caspase-1 then mediates both maturation by proteolytic cleavage and unconventional secretion of the proinflammatory cytokines interleukin-1β (IL-1β) and IL-18 (2,3). Caspase-1 also induced gasdermin D (GSDMD) cleavage, liberating its cytotoxic N-terminal fragment that oligomerizes to form large pores into the plasma membrane, leading to pyroptosis, an inflammatory cell death (4).

The NLRP3 (NOD-, LRR- and pyrin domain-containing protein 3) inflammasome (5), containing the NLRP3 PRR, is the most extensively studied inflammasome (6). Its activation requires two steps. The first, non-essential step, is priming by Toll-like receptor (TLR) agonists like lipopolysaccharide (LPS), among others, allowing transcription of genes encoding inflammasome components. The second is the activation by various infectious and stress-associated signals, with nigericin being the most potent *in vitro* (2). Activation leads to a potassium efflux out of the cell, which is the only common mechanism identified for canonical NLRP3 inflammasome activation by all stimuli (7,8). Additionally, a non-canonical pathway is triggered by intracellular LPS from phagocytosed Gram-negative bacteria that is recognized by Caspase-11 in mice or Caspases-4/5 in humans (9,10). Caspases-4/11 cleave the pore-forming protein GSDMD causing potassium efflux and NLRP3 inflammasome activation (6).

A broad range of proteins associated to different molecular mechanisms tightly governs inflammasome activation (11), including NEK7 (never in mitosis A (NIMA)-related kinase 7) (12,13). Downstream of the potassium efflux, the catalytic domain of NEK7 interacts with both the leucine-rich repeat (LRR) domain and the NACHT domain of NLRP3 (14). The formation of a NEK7-NLRP3 complex allows NLRP3-ASC complex formation, ASC oligomerization, and Caspase-1 activation (15).

The proliferating cell nuclear antigen (PCNA) is a nuclear protein essential for DNA replication and repair (16). PCNA has no enzymatic activity but is a scaffolding protein that binds to numerous proteins. In neutrophils, differentiated non-proliferative cells, PCNA is exclusively cytosolic and exerts various regulatory functions through proteins interactions. PCNA plays an anti-apoptotic role in neutrophils by interacting with pro-Caspases-3/8/9/10, preventing their activation (17). The p21 peptide and the chemical T2AA (T2 Amino Alcohol) both disrupt these interactions by binding to PCNA. They are pro-apoptotic and anti-inflammatory in different models, making PCNA a therapeutic target in inflammatory diseases (18).

Inflammasomes are mainly expressed in myeloid cells, including monocytes/macrophages and neutrophils, with molecular signaling differing between these cells (19). Since cytoplasmic PCNA regulates neutrophil survival and activation through its scaffolding function, we hypothesize that cytoplasmic PCNA could serve as a platform for regulating the NLRP3 inflammasome. This could occur for example via its interaction with pro-Caspases or other proteins such as ASC, which have homology domains (6). We investigated the NLRP3 inflammasome in monocytes/macrophages, in which the signaling pathways are clearly defined (11).

We showed that monocytes differentiation into macrophages induced a predominantly cytoplasmic relocation of PCNA. Cytoplasmic PCNA could regulate both canonical and non-canonical NLRP3 inflammasomes through its scaffolding function. Moreover, NLRP3 inflammasome stimulation caused NEK7 and potentially NLRP3 binding to cytoplasmic PCNA, and disruption of this binding impaired inflammasome activation. Finally, we suggested that the cyclin-dependent kinase inhibitor p21 might naturally repress the NLRP3 inflammasome *in vivo*. Thus, we identified PCNA as a potential new regulator of the NLRP3 inflammasome in macrophages.

## 2 MATERIALS AND METHODS

### 2.1 Cell culture

THP-1 were grown in RPMI 1640 with GlutaMAX (Gibco) supplemented with 10% FBS (Biosera), 10 mM HEPES (Gibco), 1 mM Sodium Pyruvate (Gibco), 50 µM 2-mercaptoethanol (Life Technologies), 100 U/ml penicillin and 100 μg/ml streptomycin mix (Gibco) at 37°C in a 5% CO_2_ atmosphere.

For THP-1 differentiation into macrophages, cells were cultured in 24-well plates for 3 h with 300 ng/ml Phorbol 12-myristate 13-acetate (PMA) at 3 × 10^5^ cells/well and then 3 days in complete RPMI 1640 medium without PMA. Adherent cells were used as THP-1 macrophages.

### 2.2 Reagents

Nigericin, *E. coli* LPS (0111:B4) and PMA were from Sigma-Aldrich. Monosodium urate (MSU) crystals were kindly donated by H.K. EA (INSERM UMRS1132, Université Paris Cité). All the antibodies used are listed in **Supplementary Table 1**. Hoechst 33342 solution was from ThermoScientific. µMACS Protein A Microbeads 130-071-001), MACS Separation µ Columns 130-042-701) and μMACS™ Separator were from Miltenyi Biotec.

### 2.3 Cell stimulation and PCNA inhibition

PMA-differentiated or monocytic THP-1 cells (3 × 10^5^ cells/well) were stimulated in RPMI 1640 with 10% FBS in 24-well plates. Monocytic THP-1 cells were primed with 200 ng/ml LPS during 4 h. If not otherwise indicated, NLRP3 activation was achieved with 20 µM nigericin for 1 h, 200 ng/ml MSU for 3 h or live *E. coli* strain O127: H6 (EPEC, E2348/69, kind gift from Julie Guignot) at a multiplicity of infection of 20 for 2.5 h followed by 16 h with 100 units/ml penicillin/100 μg/ml streptomycin.

To inhibit PCNA interactions, cells were pre-treated for 1 h with the p21 peptide (p21-YIRS peptide: KRRQTSMTDFYHSKRRLIFSRYIRS, Genecust) or the chemical T2AA (Sigma_Aldrich, SML0794) at 1, 5, 10 or 25 µM before NLRP3 activation by nigericin or MSU. The mutated homologue p21 peptide (p21m) (p21 mute-YIRS peptide: KRRQTGETDFDHAKAALIFSRYIRS, Genecust), not allowing inhibition, was used as control.

For *E. coli*. infection, p21, p21m or T2AA were added concomitantly with penicillin/streptomycin. After stimulation, culture supernatants were collected, centrifuged 5 min at 2000 rpm and stored at −20°C. For each experiment, each culture condition was performed in triplicate.

### 2.4 ELISA

To detect secreted IL-1β, collected supernatants were mixed with PBS-Tween 20 (0.05% final concentration) and heated for 5 min at 95°C, as previously described (20). IL-1β levels were measured using human (Invitrogen, 88-7261-88) or mouse (Invitrogen, 88-7013-88) IL-1 beta Uncoated ELISA kit according to the manufacturer’s instructions.

### 2.5 Immunofluorescence labeling and microscopy analysis

To perform cytospins, 2 x 10^5^ THP-1 cells or mouse BMDMs were cytocentrifuged on SuperFrostTM Plus adhesion slides (Epredia) 3 min at 300 g using Cytospin 3 (Shandon). Cells were fixed in PBS-4% formaldehyde (Sigma) for 20 min on ice, permeabilized with 0.25% Triton X-100 for 5 min at RT, and blocked with PBS-1% BSA for 30 min at RT. Cells were incubated at RT with PBS 1% BSA-diluted primary antibodies (PCNA ab5, 1/25) for 1 h followed by biotinylated antibodies (1:150) for 30 min and streptavidin-coupled Alexa Fluor 555 (1:1000) for 30 min. Nuclei were stained with Hoechst 33342 (1:6000 in PBS) for 10 min. Slides were mounted using Fluoromount-G^TM^ medium (Invitrogen). Image were acquired at the Cochin Institute Imaging Facility (IMAG’IC) using the Widefield Zeiss Observer Z1. Digital images were analyzed using Inscoper_7.2.1 software (Molecular Devices) and quantified using the ImageJ 1.53e software.

### 2.6 Duolink® Proximity Ligation Assay (PLA)

Duolink II proximity ligation assay kit (PLA-probe anti-rabbit plus, catalog #DUO92004-30RXN; PLA-probe anti-mouse minus, catalog #DUO92006-30RXN; Detection Kit Orange, catalog #DUO92007-100RXN) was used following the manufacturer’s instructions (Merck Sigma-Aldrich). Cytospsin slides of THP-1 cells were prepared as described for immunofluorescence staining. Anti-PCNA (ab5; 1:200), anti-ASC (1:200) and anti-NEK7 (1:50) were used as primary antibodies. Slides were mounted with the Duolink in situ mounting medium with DAPI (Sigma). Cells were observed with a Widefield Zeiss Observer Z1. The fluorescent signal is generated only if the proteins of interest are located within 40 nm. Digital images were analyzed and quantified as described for immunofluorescence staining.

### 2.7 Co-immunoprecipitation (co-IP)

THP-1 macrophages were lysed by sonication in 50 mM HEPES, 4 mM PMSF, 400 µM leupeptin, 400 µM pepstatin, 1 mM orthovanadate, 1 mM EGTA, 1 mM EDTA and 1 mM DTT, during 10 seconds on ice (Soniprep 150 plus, MSE) and centrifuged 10 min at 14 000 rpm at 4°C to obtain cytosolic contents. Protein concentration was determined using BCA assay (Pierce). Cytosolic proteins (500 µg) were incubated with 10 µl of protein A microbeads for 1 h at 4°C under rotation. The lysate was passed through a µ Columns in a magnetic field of the μMACS™ Separator to recover a pre-cleared lysate. The lysate was then incubated with 1 µg anti-PCNA pAb (ThermoFisher) or control isotype Ab (Cell Signaling) for 1 h at 4°C under rotation. Then, 50 µl protein A microbeads were added and incubated for 1 h at 4°C under rotation. This mix was loaded onto a column containing Sepharose-coated magnetic beads and washed four times with a stringent washing buffer (50 mM Tris, 300 mM NaCl and 1% NP-40, pH 8), followed by five washes with a hyposaline buffer (20 mM Tris, pH 7.5). Proteins were eluted with 50 µl of preheated 5× Laemmli buffer and analyzed by Western blot.

### 2.8 Western blot analysis

Total proteins were extracted using lysis buffer (10 mM Tris–HCl, 150 mM NaCl pH 7.8, 1% Nonidet P-40 (Sigma-Aldrich), protease inhibitors cocktail (complete mini EDTA-free, Roche Diagnostics)). Alternatively, cells were washed with PBS, lysed with 5x Laemmli buffer and sonicated (5 cycles). Protein concentration was determined using BCA assay (Pierce). 30 µg of proteins were fractionated on SDS-PAGE and transferred to a polyvinylidene difluoride membrane (Millipore). The membrane was blocked in Tris-buffered saline (TBS) containing 5% non-fat dry milk for 1 h at RT and probed with primary antibodies at 4°C overnight (PCNA PC10, 1:1000). β-actin or tubulin were used as loading control. Horseradish peroxidase-conjugated secondary antibodies diluted in TBS 5% non-fat dry milk were incubated 1 h at RT, followed by enhanced chemiluminescence detection (Amersham^TM^, Cytiva). Images were acquired using the Fusion FX7 system (Vilbert Lourmat).

### 2.9 Statistical analysis

Statistical analysis was performed using the R software. IL-1β secretion was measured in several independent experiments and normalized to the mean value of the positive control in each experiment. For western blot quantification, data were normalized to the positive control in each experiment. Comparisons of all the normalized values were made using the non-parametric Mann-Whitney test. Differences were considered significant when P < 0.05.

## 3 RESULTS

### 3.1 Differentiation of THP-1 monocytes into macrophages induces cytoplasmic relocation of PCNA

THP-1 human myelomonocytic cell line is commonly used for inflammasome research. PCNA was localized in both nucleus and cytoplasm of THP-1 monocytes as attested by immunofluorescence staining using 3 different anti-PCNA antibodies (**Figure 1A and Supplementary Figure S1A**). However, PCNA became predominantly cytoplasmic after differentiation into macrophages by PMA treatment (**Figure 1A and Supplementary Figure S1A**). Confirming these results, western blot analysis showed that PCNA protein was expressed in equal amount in the cytoplasm and nucleus of THP-1 monocytes, but was mainly expressed in the cytoplasm of THP-1 macrophages (**Figure 1B, C**). Expression of lamin exclusively in the nucleus confirmed that the cytoplasmic fraction was not contaminated by nuclear proteins, allowing us to conclude that PCNA expression in the cytoplasm was the consequence of the relocation of the protein from the nucleus to the cytoplasm, and not of contamination by nuclear proteins (**Figure 1B**). As previously described (20), NLRP3 inflammasome activation by nigericin induced IL-1β secretion by both LPS-primed THP-1 monocytes and PMA-differentiated and PMA-primed THP-1 macrophages (**Supplementary Figures S1B, C**). Since NLRP3 inflammasome is a cytoplasmic complex, and since THP-1 macrophages expressed very few nuclear PCNA, this suggested that if PCNA was necessary, it is primarily the cytoplasmic form that should be required for NLRP3 inflammasome activation by nigericin. Moreover, nigericin treatment did not affect the amount of PCNA protein expressed in the cytoplasm of THP-1 macrophages **(Figure 1D, E)** indicating that no PCNA degradation occurred during nigericin treatment. By the same way, nigericin treatment did not affect the relocation of PCNA into the cytoplasm in PMA-differentiated THP-1 macrophages **(Figure 1F).**

**FIGURE 1.**
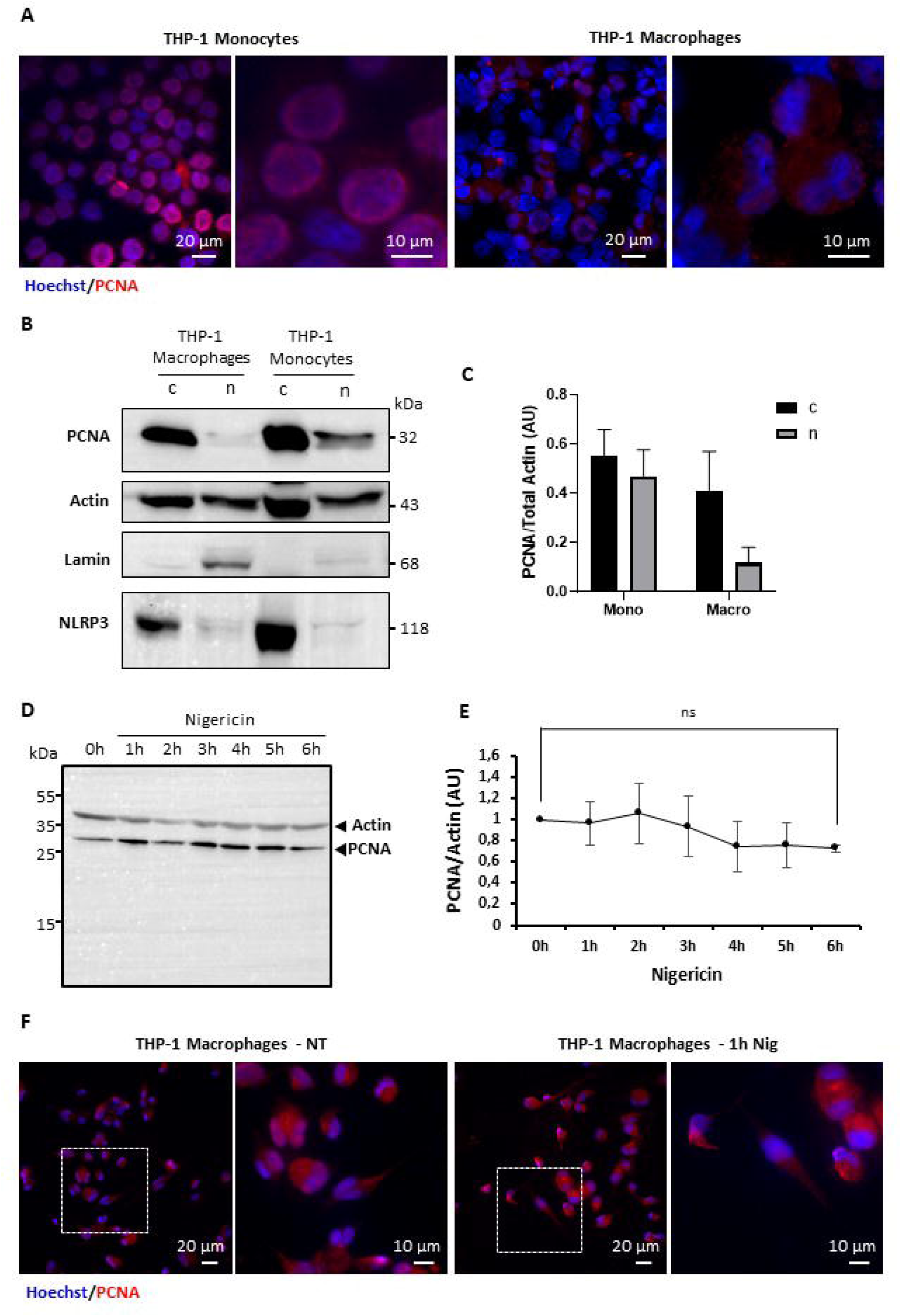
PCNA expression and canonical NLRP3 inflammasome activation in monocytic and macrophagic human THP-1 cells. **(A)** Localization of PCNA using PCNA-specific immunofluorescence microscopy analysis performed on monocytic THP-1 cell line or THP-1 cells differentiated into macrophages by PMA (300 ng/ml for 3 h followed by 3 days of resting). Red, PCNA; blue, Hoechst. Representative micrographs are shown. **(B)** Western blot analysis of PCNA expressed in nuclear (n) and cytoplasmic (c) protein lysates from PMA-differentiated THP-1 macrophages. 20 and 40 µg nuclear and cytoplasmic proteins were submitted to electrophoresis, respectively. β-actin, lamin and NLRP3 serve as total, nuclear and cytoplasmic loading control, respectively. **(C)** Densitometric analysis of bands obtained in (B). Cytoplasmic (c) and nuclear (n) PCNA were obtained by establishing the PCNA/Total Actin (c actin + n actin) ratios, related to the same amount of protein. AU, arbitrary units. **(D)** Representative western blot analysis of PCNA expressed in protein lysates from PMA-differentiated THP-1 macrophages untreated (0h) or treated by nigericin during the indicated time (h, hours). β-actin serves as control. Representative of three independent experiments. **(E)** Densitometric analysis of PCNA and β-actin bands obtained in western blot analysis of protein lysates from PMA-differentiated THP-1 macrophages untreated (0h) or treated by nigericin during the indicated time (h, hours). n=3 (Figure 1D and not shown) except for 6h treatment where n=2. Data represents mean PCNA/Actin ratios ± sem (AU, arbitrary units). NS, not significant. **(F)** PCNA-specific immunofluorescence microscopy analysis of PMA-differentiated THP-1 macrophages treated (1 h Nig) or not (NT) with 20 µM nigericin during 1 h. Red, PCNA; blue, Hoechst. Representative micrographs are shown. Surrounded cells are magnified.

### 3.2 Preventing PCNA partners binding inhibits activation of the canonical and non-canonical NLRP3 inflammasomes in THP-1 macrophages

We tested whether preventing PCNA scaffold could impair NLRP3 inflammasome activation. We used the p21 peptide and the T2AA, both of which dissociate PCNA interactions by binding to it. Neither compound inhibited IL-1β secretion induced by LPS priming and nigericin treatment in THP-1 monocytes (**Figure 2A and Supplementary Figure S2A**). However, in PMA-differentiated THP-1 macrophages, in which PCNA was mostly cytoplasmic, the p21 peptide significantly inhibited nigericin-induced IL-1β secretion, unlike the mutant p21 peptide deprived of PCNA binding sites or the T2AA (**Figure 2B and Supplementary Figure S2B**). As expected, NLRP3 inflammasome activation by nigericin induced cleavage of both pro-Caspase-1 and GSDMD in THP-1 macrophages (**Figure 2C**). Consistent with the IL-1β secretion observed, the p21 peptide, but not the mutant p21 peptide, significantly inhibited the cleavage of both pro-Caspase-1 and GSDMD (**Figure 2C**), whereas none of these treatments significantly affected PCNA expression (**Figure 2D**).

**FIGURE 2.**
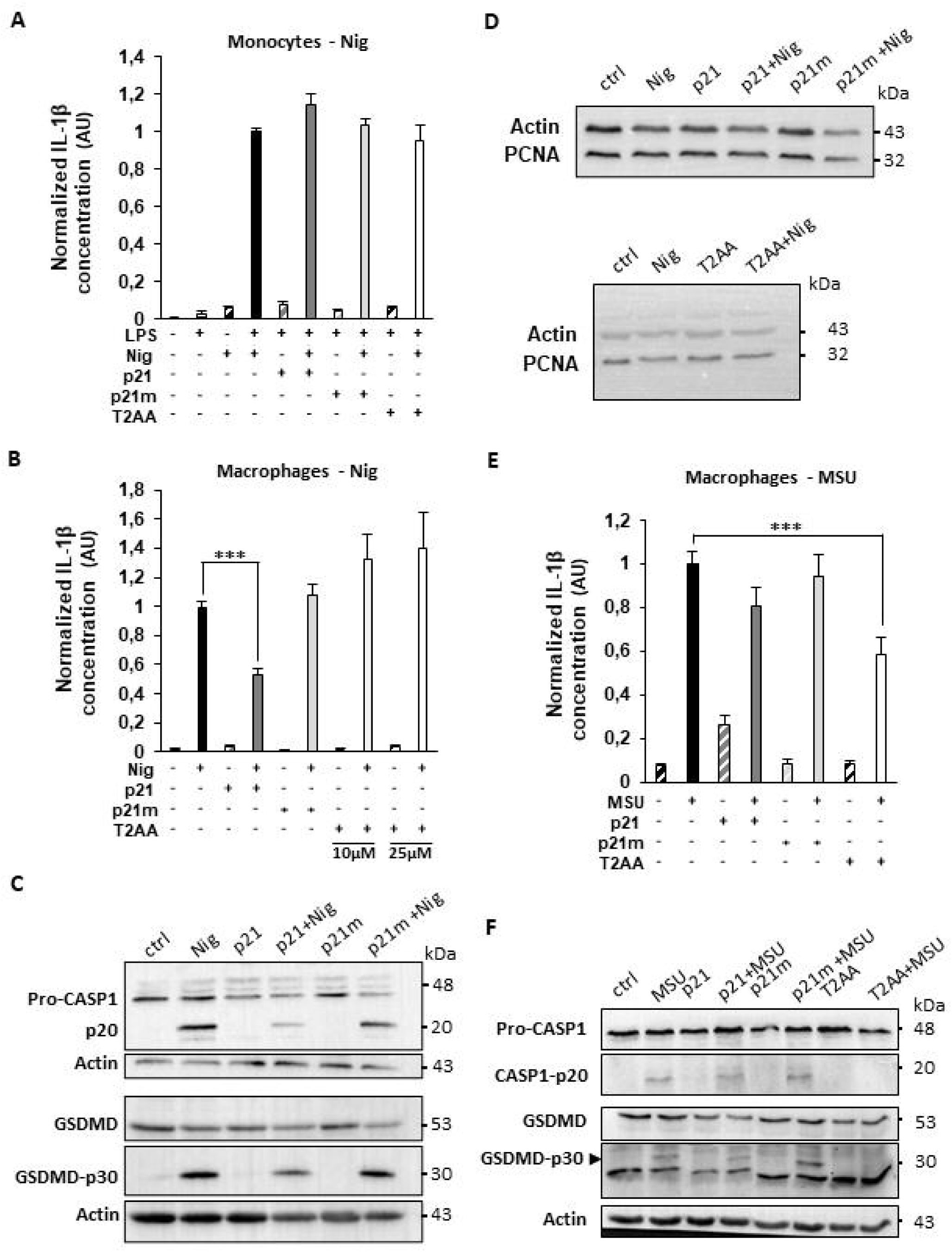
Preventing the interaction of PCNA with its partners by the p21 peptide or the T2AA inhibits NLRP3 canonical inflammasome activation in THP-1 macrophagic cell line. **(A, B)** ELISA quantification of IL-1β secreted by **(A)** THP-1 monocytes (6 independent experiments performed in duplicate (n=12) or triplicate (n=14), depending the conditions) cultured for 4 h with (+) or without (−) 200 ng/ml LPS pretreated (+) or not (-) with 10 µM p21 peptide (p21) or mutant p21 peptide (p21m) or 25 µM T2AA, and **(B)** PMA-differentiated THP-1 macrophages pretreated (+) or not (-) (16 independent experiments performed in single sample (n=1), duplicate (n=12), triplicate (n=2) or quadruplicate (n=1); total n=35) with 10 µM p21 (13 independent experiments performed in duplicate (n=12) or quadruplicate (n=1); total n=28) or p21m (9 independent experiments performed in duplicate (n=8) or quadruplicate (n=1); total n=20), 10 µM T2AA (light grey histograms, 11 independent experiments performed in duplicate (n=6), triplicate (n=4) or quadruplicate (n=1); total n=28) or 25 µM T2AA (white histograms, 7 independent experiments performed in duplicate (n=3) or triplicate (n=4); total n=18) during 1 h, followed by incubation in absence (−) or presence (+) of 20 µM nigericin (Nig) during 1 h. **(C)** Western blot analysis of CASPASE-1 (Pro-CASP1, pro-CASPASE-1; p20, cleaved form of CASPASE-1) and GASDERMIN D (p30, N-terminal cleaved fragment of GSDMD) in PMA-differentiated THP-1 macrophages untreated (ctrl) or pretreated 1 h with or without 10 µM p21 or 10 µM p21m followed by 1 h incubation with or without 20 µM nigericin. β-actin serves as a loading control. Representative of three independent experiments. **(D)** Representative western blot analysis of PCNA in PMA-differentiated THP-1 macrophages untreated (ctrl) or pretreated 1 h with or without 10 µM p21 or 10 µM p21m or 10 μM T2AA followed by 1 h incubation with or without 20 µM nigericin. β-actin serves as a loading control. **(E)** ELISA quantification of IL-1β secreted by PMA-differentiated THP-1 macrophages pretreated 1h in absence (−) (7 independent experiments performed in single sample (n=2), duplicate (n=3) or triplicate (n=2); total n=14) or presence (+) of 10 µM p21 (7 independent experiments performed in single sample (n=2), duplicate (n=3) or triplicate (n=2); total n=14), 10 µM p21m (7 independent experiments performed in duplicate (n=6) or triplicate (n=1); total n=15) or 25 µM T2AA (7 independent experiments performed in single sample (n=1), duplicate (n=4) or triplicate (n=2); total n=15) then cultured for 3 h with (+) or without (−) 200 µg/ml monosodium urate (MSU) crystals. **(F)** Western blot analysis of CASPASE-1 (Pro-CASP1, pro-CASPASE-1; p20, cleaved form of CASPASE-1) and GASDERMIN D (p30, N-terminal cleaved fragment of GSDMD) in PMA-differentiated THP-1 macrophages untreated (ctrl) or pretreated 1 h with or without 10 µM p21, 10 µM p21m or 25 µM T2AA followed by 1 h incubation with or without 200 µg/ml MSU crystals. β-actin serves as a loading control. For GSDMD, representative of three independent experiments. (A, B, E) Results are expressed as mean ± sem of normalized data from independent experiments (AU, arbitrary units). For each independent experiment, results were normalized to the mean of the positive control (LPS+ Nig+ for THP-1 monocytes (ranging from 995 to 6551 pg/ml) (**Supplementary Figure S2A**); Nig+ for THP-1 macrophages (ranging from 312 to 12178 pg/ml) (**Supplementary Figure S2B**); MSU+ for THP-1 macrophages (ranging from 313 to 2045 pg/ml) (**Supplementary Figure S2C**)). Non-parametric Mann-Whitney tests were performed, ***p < 0.001.

MSU crystals, which are NLRP3 inflammasome activators implicated in gout disease (21), induced IL-1β secretion by PMA-differentiated THP-1 macrophages (**Figure 2E and Supplementary Figure S2C**). Although p21 induced a slight decrease in IL-1β secretion, T2AA significantly decreased it, along with inhibition of pro-Caspase-1 and GSDMD cleavage (**Figure 2E, F**).

To investigate whether cytoplasmic PCNA could also regulate the non-canonical NLRP3 inflammasome, PMA-differentiated THP-1 macrophages were cultured with enteropathogenic *Escherichia coli* (EPEC) gram-negative bacteria. As expected, EPEC induced a strong IL-1β secretion by THP-1 macrophages, which was significantly inhibited by both p21 and T2AA, but not by p21m (**Figure 3A and Supplementary Figure S2D**). As previously described (10), upon activation by intracellular LPS from phagocytosed bacteria, Caspase-4 cleaves GSDMD to induce pyroptosis (**Figure 3B**), leading to consequent activation of the Caspase-1-dependent-NLRP3 inflammasome and IL-1β secretion. Whereas IL-1β secretion was decreased in presence of p21 and T2AA (**Figure 3A**), none of these inhibitors diminished the Caspase-4-mediated GSDMD cleavage in THP-1 macrophages (**Figure 3B**). These results suggested that cytoplasmic PCNA may regulate the non-canonical NLRP3 inflammasome through a mechanism not involving the Caspase-4/GSDMD axis.

**FIGURE 3.**
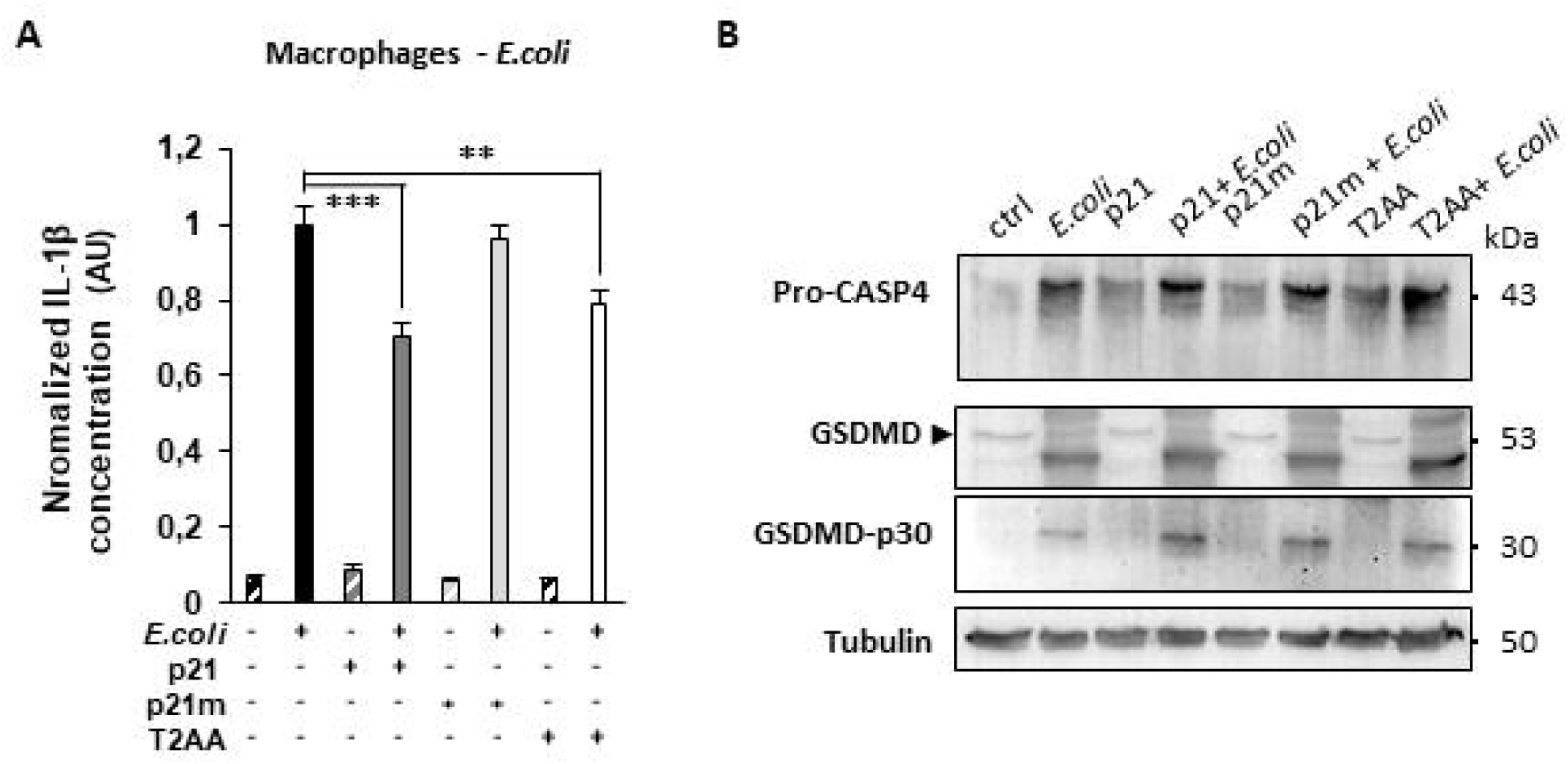
Preventing the interaction of PCNA with its partners by the p21 peptide or the T2AA inhibits NLRP3 non-canonical inflammasome activation in THP-1 macrophagic cell line. **(A)** ELISA quantification of IL-1β secreted by PMA-differentiated THP-1 macrophages cultured for 2,5 h with (+) or without (−) *E. coli* bacteria at a multiplicity of infection of 20, followed by a 16 h incubation period with penicillin/streptomycin in absence (−) or presence (+) of 5 µM p21, 5 µM p21m or 10 µM T2AA (4 independent experiments performed in triplicate or quadruplicate, depending the conditions; total n=12 to 14). Results are expressed as mean ± sem of normalized data from independent experiments (AU, arbitrary units). For each independent experiment, results were normalized to the mean of the positive control (*E. coli*+ ranging from 1705 to 8366 pg/ml (**Supplementary Figure S2D**)). Non-parametric Mann-Whitney tests were performed, **p < 0.01; ***p < 0.001. **(B)** Western blot analysis of CASPASE-4 (Pro-CASP4, pro-CASPASE-4) and GASDERMIN D (p30, N-terminal cleaved fragment of GSDMD) in PMA-differentiated THP-1 macrophages cultured for 2,5 h with (+*E. coli*) or without *E. coli* bacteria at a multiplicity of infection of 20, followed by a 16 h incubation period with penicillin/streptomycin in absence or presence of 5 µM p21, 5 µM p21m or 10 µM T2AA. Tubulin serves as a loading control. Caspase-4 antibody detects pro-CASP4 (45 kDa) and processing intermediate forms of CASP4 (40 kDa and 32 kDa). For GSDMD, representative of two independent experiments.

### 3.3 The scaffolding protein PCNA could regulate the NLRP3 inflammasome by interacting with NEK7

To elucidate the molecular mechanisms by which the cytoplasmic PCNA could control the NLRP3 inflammasome in THP-1 macrophages, proximity ligation assay was performed. PCNA and ASC showed minimal association in resting THP-1 macrophages, but moved progressively closer together following NLRP3 activation by nigericin (**Figure 4A**). However, PCNA protein co-immunoprecipitation assay performed on cytosolic proteins obtained by sonication treatment suggested no interaction between cytosolic PCNA and ASC whatever the activation state (**Figure 4B**). In contrary, preliminary data suggested that cytosolic PCNA might associate with the NLRP3 sensor both before and after nigericin stimulation (**Supplementary Figure S3A**). The colocalization of NLRP3 and PCNA near the nucleus (**Supplementary Figure S3B**) correlated with speck formation during inflammasome activation. Moreover, nigericin treatment could led to cytoplasmic PCNA interaction with the kinase NEK7 (**Figure 4C-F**). PLA showed that 18% of resting THP-1 macrophages showed PCNA-NEK7 proximity, predominantly in the nucleus, possibly explaining why NEK7 wasn’t detected in cytosolic protein co-immunoprecipitation. After nigericin stimulation, 28% of THP-1 macrophages presented PCNA-NEK7 interaction, primarily in the cytoplasm, consistent with the inflammasome complex’s cytoplasmic location (**Figure 4C, D**). Finally, cells pretreatment with p21 seemed to strongly decrease the interaction between cytoplasmic PCNA and NEK7 (**Figure 4F**) and NLRP3 (**Supplementary Figure S3A**). These data strongly suggested that cytoplasmic PCNA may belong to the NLRP3 inflammasome complex by interacting with NEK7 and potentially NLRP3 in THP-1 macrophages.

**FIGURE 4.**
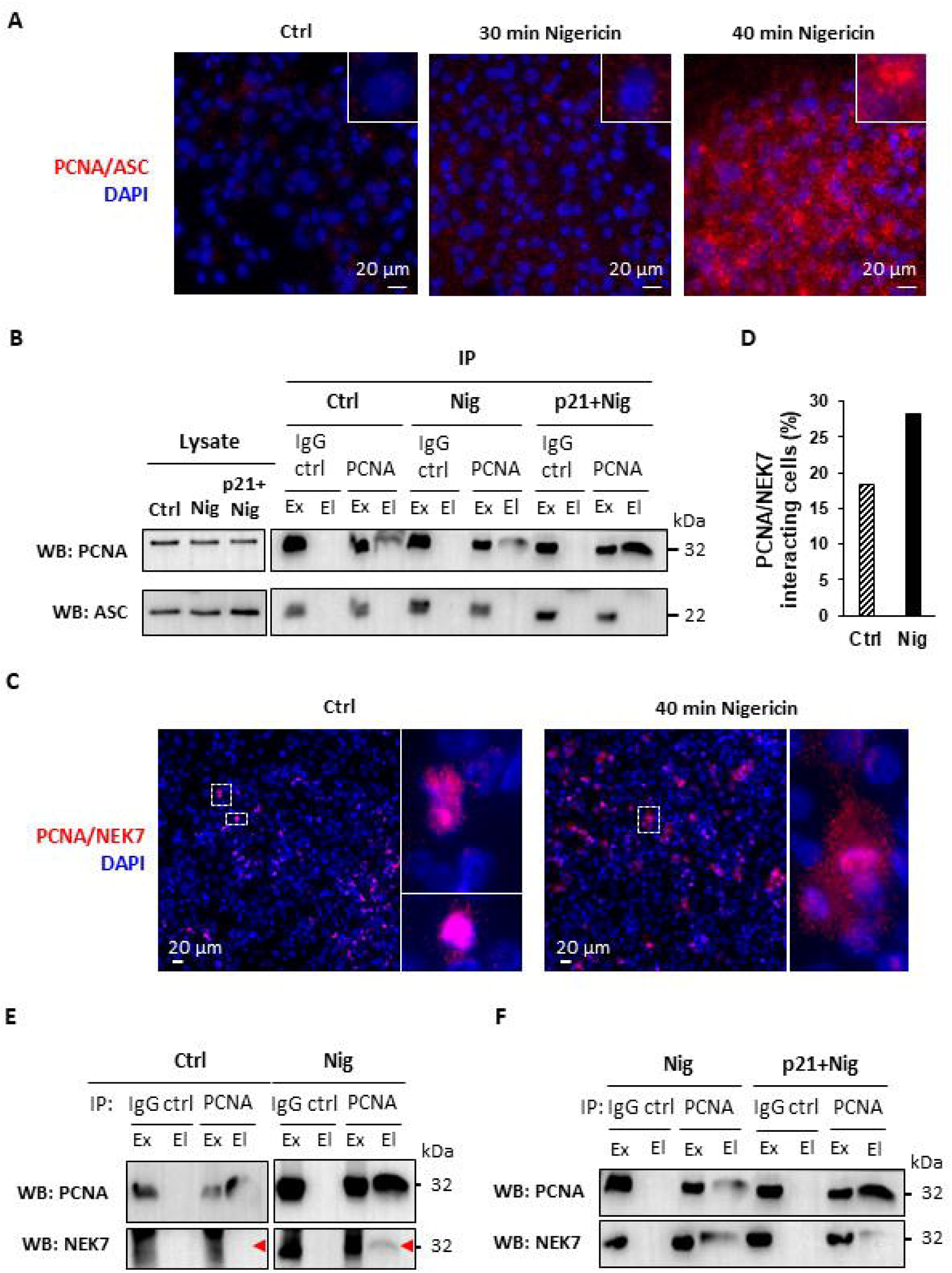
PCNA could be a scaffold for NLRP3-NEK7 interaction. **(A)** Assessment of the PCNA-ASC interaction in PMA-differentiated THP-1 macrophages using the Duolink**®** proximity ligation assay (n=1). Cells were untreated (ctrl) or treated with 20 µM nigericin during the indicated times. PCNA-ASC proximity induced a red fluorescence. DAPI was used for nuclear staining (blue). Surrounded cells are magnified 3 times. **(B)** Co-immunoprecipitation (co-IP) experiments were performed on cytosol obtained by sonication of PMA-differentiated THP-1 macrophages untreated (ctrl), treated 40 min with 20 µM nigericin (Nig) or pretreated 1 h with 10 µM p21 followed by 40 min incubation with 20 µM nigericin (p21+Nig). Co-IP using beads coated with anti-PCNA pAb (IP: PCNA) or with control isotype Ab (IP: IgG ctrl) were performed. Indicated proteins were detected by western blot analysis (WB) on the exclusion (Ex) and elution (El) fractions obtained by co-IP. For ASC detection, representative of three independent experiments (except p21+Nig, n=1). **(C)** Assessment of the PCNA-NEK7 interaction in PMA-differentiated THP-1 macrophages using the Duolink® proximity ligation assay. Cells were untreated (ctrl) or treated with 20 μM nigericin during 40 min. PCNA-NEK7 proximity induced a red/pink fluorescence. DAPI was used for nuclear staining (blue). Surrounded cells are magnified 7 times. **(D)** Quantification of the number of cells with a PCNA-NEK7 proximity-induced fluorescence in (C). A total of 694 and 823 cells for untreated (ctrl) and nigericin-treated (Nig) PMA-differentiated THP-1 macrophages, respectively, was counted. Results are expressed as the percentage of PCNA-NEK7 proximity-induced fluorescence positive cells. **(E, F)** Co-IP experiments were performed as in (B).

## 4 DISCUSSION

Tight regulation of the NLRP3 inflammasome is critical to prevent an involuntary or uncontrolled inflammatory response. Numerous mechanisms are involved in this regulation (22). Here we highlighted the interaction between cytoplasmic PCNA and its binding partners as a novel regulator of the NLRP3 inflammasome activation. Indeed, disruption of cytoplasmic PCNA interactions impaired NLRP3 inflammasome activation in THP-1 macrophages, reducing IL-1 β secretion, Caspase-1 activation and GSDMD cleavage. Since NLRP3 inflammasome assembly occurs in the cytoplasm, cytoplasmic localization of PCNA may be a prerequisite to assume its inflammasome regulatory function. Indeed, we showed that in THP-1 monocytes, where PCNA was mainly nuclear, neither p21 nor T2AA was able to inhibit IL-1 β secretion. Since PCNA is exclusively localized in the cytoplasm of neutrophils, where it was initially described to interact with apoptotic Caspases (17), it will be interesting to evaluate if PCNA could regulate the NLRP3 inflammasome in these cells. More generally, the potential inflammasome regulatory function of PCNA should be tested in mature differentiated cells in which cytoplasmic relocation of PCNA has occurred. The p21 peptide is derived from the p21/waf1/CIP1 protein, a cyclin-dependent kinase inhibitor that binds to nuclear PCNA to regulate its chromatin-bound protein functions (23). Bone marrow derived-macrophages (BMDMs) from wild type (WT) and p21/waf1/CIP1 knockout (p21 KO) mice expressed PCNA mainly in the cytoplasm. Our preliminary data showed that p21 KO BMDMs could secrete higher IL-1β levels than WT BMDMs following NLRP3 inflammasome stimulation (**Supplementary Figure S4)**. These results suggested that p21 protein might be a natural NLRP3 inflammasome repressor.

In our study, both canonical and non-canonical NLRP3 inflammasomes activation was inhibited by agents that disrupted cytoplasmic PCNA interactions. In the canonical pathway, p21 and/or T2AA prevented Caspase-1 activation, thus inhibiting GSDMD Cleavage and IL-1β secretion. In the non-canonical pathway, we showed that neither cleavage of pro-Caspase-4 nor the consequent GSDMD cleavage was affected by these two inhibitors, whereas IL-1β secretion was decreased. These results suggested that PCNA should not interfere with the Caspase-4/GSDMD axis of the non-canonical pathway and that cytoplasmic PCNA may regulate the inflammasome by acting on the canonical part of the complex (Caspase-1 axis). Neither THP-1 cells nor mouse BMDMs expressed an alternative NLRP3 inflammasome (24), precluding investigation of the cytoplasmic PCNA role in this pathway. Alternative inflammasome induced in human monocytes/macrophages by extracellular LPS involves a RIPK1-FADD-Caspase-8 complex triggering the NLRP3-ASC-Caspase-1 complex (24,25). It will be interesting to test whether cytoplasmic PCNA is able to impair the NLRP3 alternative inflammasome activation by targeting Caspase-1 activation. Of note, Caspase-8 is involved in the canonical, non-canonical and alternative NLRP3 inflammasomes (24,26). We can hypothesize that cytoplasmic PCNA might regulate inflammasome activation regulating Caspase-8 availability, given that PCNA interaction with Caspase-8 promotes cell survival in neutrophils (17). Thus, it is possible that Caspase-8 could also partner with cytoplasmic PCNA during NLRP3 inflammasome activation.

The p21 peptide has the highest known interaction affinity for PCNA (23), thus acting as a competitive inhibitor for the binding of PCNA partners. Most of the PCNA-interacting proteins, including p21, possesses a short amino acid sequence, called a PIP-box, binding to the interdomain connecting loop (IDCL) of PCNA (27). T2AA, a non-peptide inhibitor derived from T3 (3,3′,5-triiodothyronine) but lacking thyroid hormone activity, also binds to the IDCL and to a lesser extent to a shallow cavity present at the PCNA homotrimer interface adjacent to lysin 164 (28). T2AA disrupts interactions involving monoubiquitinated PCNA and its partners. Depending the stimulus, either p21, T2AA or both were able to inhibit inflammasome activation in THP-1 macrophages, suggesting that different proteins could intervene to regulate NLRP3 inflammasome activation. These proteins probably bind different sites on PCNA, explaining why p21 and T2AA were not always able to inhibit inflammasome activation. Moreover, it is possible that different forms of PCNA may be implicated (trimeric, monomeric, ubiquitinated…) according to the stimulus used. It should be noted that NEK7 and NLRP3, the two proteins we found potentially interacting with PCNA, lack a PIP-box. This suggests they may bind to the IDCL through alternative motif (29), or to other sites on PCNA, or indirectly through interaction with other proteins. In our case, we can hypothesize that NEK7 may be indirectly linked to PCNA through at least interaction with NLRP3. Further studies are required to identify the binding sites of PCNA with NEK7 and/or potentially NLRP3, and to identify potential other proteins implicating in NLRP3 inflammasome complex activation and interacting with cytoplasmic PCNA.

In resting cells, the NLRP3 inflammasome is kept inactive to prevent unwanted inflammation (30). NLRP3 is sequestered in a double ring cage structure associated with organelles membranes (31–33). Since our results suggested that cytoplasmic PCNA and NLRP3 could interact in resting THP-1 macrophages, and considering the ring shape structure of trimeric PCNA (27), we can hypothesize that cytoplasmic PCNA could contribute to NLRP3 caging. Following inflammasome stimulation, NLRP3 should travel from the membranes to the centrosome/microtubule-organizing center (MTOC), to interact with NEK7 located there. Previous report had shown that PCNA can bind γ-tubulin that expresses a PIP motif, facilitating PCNA transport along microtubules from cytosol to nucleus (34). We can hypothesize a similar PCNA-tubulin interaction may transport the potential PCNA-NLRP3 complex from membranes to the MTOC. At the MTOC, NLRP3-NEK7 interaction disrupts the NLRP3 cage structure (30–32). We showed that PCNA and NEK7 colocalized in the nucleus in resting THP-1 macrophages, but in the cytoplasm after nigericin treatment. Moreover, we found that after nigericin treatment, PCNA interacted with NEK7 and potentially NLRP3. These results suggested that PCNA, through its tubulin interaction, might also participate in the transport of NEK7 from the nucleus to the cytoplasm. Only one study previously described a PCNA-NEK7 interaction, and it was not in the context of inflammasome activation (35). We propose that cytoplasmic PCNA-NEK7 binding at the MTOC could facilitate the NLRP3 cage disruption and subsequent NLRP3-NEK7 interaction, allowing ASC oligomerization. The observation that ASC and PCNA moved closer together after inflammasome stimulation, without direct interaction, is in agreement with our previous hypotheses. Furthermore, since p21 peptide partially disrupted the PCNA-NEK7 interaction and since p21/waf1/CIP1 KO BMDMs could secrete higher IL-1β levels following NLRP3 inflammasome stimulation, suggested that p21 protein might be a natural inflammasome repressor, preventing PCNA-NEK7 interaction.

In summary, our study highlighted cytoplasmic PCNA as a potential novel component of the NLRP3 inflammasome acting as a regulator of inflammasome assembly, and thus of the subsequent Caspase-1 activation, GSDMD cleavage and IL-1β secretion. Our study will undoubtedly pave the way for future research aimed at exploring the role of cytoplasmic PCNA in different types of inflammasomes and at identifying its different partners and their binding site depending on the activation stimulus.

## AUTHOR CONTRIBUTIONS

LT conceived, designed and supervised the study. LT, CL and KA designed and performed the experiments. LT and KA managed the breeding of mouse colony. CL, KA and LT analyzed the results. NC performed the statistical analyses. LT and CL wrote the manuscript. LT and GF performed the revision of the manuscript. All authors reviewed and accepted the final version of the manuscript.

## ACKNOWLEDGMENTS

We thank H.K. Ea (INSERM UMRS1132, Université Paris-Cité, hôpital Lariboisière, Paris, France) for providing MSU crystals and J. Guignot (Cochin Institute) for providing EPEC. We thank A-S. Dangeard (Cochin Institute) for her help with handling in the bacteriology laboratory. We thank D. Judith for discussions on co-immunoprecipitation experimental design. We thank the MOUSET’IC Cochin Institute (Marcio Do Cruzeiro) core facility for his grateful help with mouse genotyping and maintenance of the line. We also thank the IMAG’IC Cochin Institute core facility.

