## Supplemental Table 1 for "The cytoplasmic scaffolding protein PCNA regulates NLRP3 inflammasome activation in macrophages"

**Supplementary Table S1.** List of antibodies used in the study

| **Antibody** | **Supplier (Cat.number)** | **Type** | **Host** | **Application** | **Dilution** |
| --- | --- | --- | --- | --- | --- |
| Actin | Sigma-Aldrich (A2066) | Polyclonal/Primary | Rabbit | WB | 1:5000 |
| Anti-mouse IgG (H+L), Biotinylated | Vector Laboratories  (BP-9200-50) | Secondary Biotinylated | Goat | IF | 1:1500 |
| Anti-Rabbit Immuglobulins/Biotinylated (E04232) | Dako (BA-200) | Secondary Biotinylated | Goat | IF | 1:250 |
| ASC/TMS1/PYCARD(B-3) | Santa Cruz  (sc-514414) | Monoclonal/Primary | Mouse | IF/PLA | 1:200 |
| β-Tubulin | Cell Signaling (2146S) | Polyclonal/  Primary | Rabbit | WB | 1:1000 |
| Caspase-1 (D7F10) | Cell Signaling (3866S) | Monoclonal/Primary | Rabbit | WB | 1:1000 |
| Caspase-4 | Cell Signaling (4450S) | Polyclonal/  Primary | Rabbit | WB | 1:1000 |
| Cleaved Caspase-1 Asp297 (D57A2) | Cell Signaling (4199S) | Monoconal/Primary | Rabbit | WB | 1:1000 |
| Cleaved GSDMD Asp275 (E7H9G) | Cell Signaling (36425S) | Monoconal/Primary | Rabbit | WB | 1:1000 |
| Cryopyrin/NALP3/NLRP3 (H-8) | Santa Cruz  (sc-518123) | Monoclonal/Primary | Mouse | IF | 1:400 |
| GSDMD | Proteintech (20770-1-AP) | Polyclonal/  Primary | Rabbit | WB | 1:1000 |
| Isotype control (DA1E) IgG XP(R) | Cell Signaling (3900S) | Monoclonal | Rabbit | CO-IP | 1µg |
| Lamin B1 (D9V6H) | Cell Signaling (13435S) | Monoconal/Primary | Rabbit | WB | 1:1000 |
| NLRP3 (D2P5E) | Cell Signaling (13158S) | Monoclonal/Primary | Rabbit | WB | 1:1000 |
| NEK7 (C34C3) | Cell Signaling (3057S) | Monoclonal/Primary | Rabbit | IF,WB | 1:50, 1:1000 |
| PCNA (PC10) | Dako (MO87) | Monoclonal/Primary | Mouse | WB, IF | 1:1000, 1:250 |
| PCNA (ab5) | Sigma-Aldrich (PC474) | Polyclonal/  Primary | Rabbit | IF, PLA, WB | 1:250, 1:200, 1:1000 |
| PCNA (1D7) | Abbkine (ABL1040) | Monoclonal/Primary | Mouse | IF | 1:100 |
| PCNA | ThermoFisher (PA5-81628) | Polyclonal/  Primary | Rabbit | CO-IP | 1µg |
| Peroxidase-conjugated AffiniPure anti-rabbit IgG, F(ab')₂ fragment specific | Jackson ImmunoResearch (111-035-006) | Polyclonal/  Secondary | Goat | WB | 1:1000 |
| Peroxidase-conjugated AffiniPure anti-mouse IgG, F(ab')₂ fragment specific | Jackson ImmunoResearch (115-035-006) | Polyclonal/  Secondary | Goat | WB | 1:1000 |
| Streptavidin Alexa Fluor 555 conjugate | Invitrogen (S32355) | Tertiary/  Streptavidin conjugate | - | IF | 1:2000 |
| TMS1/ASC (E1E3I) | Cell Signaling (13833S) | Monoclonal/Primary | Rabbit | WB | 1:1000 |
