## Supplemental Figure 1 for "The cytoplasmic scaffolding protein PCNA regulates NLRP3 inflammasome activation in macrophages"

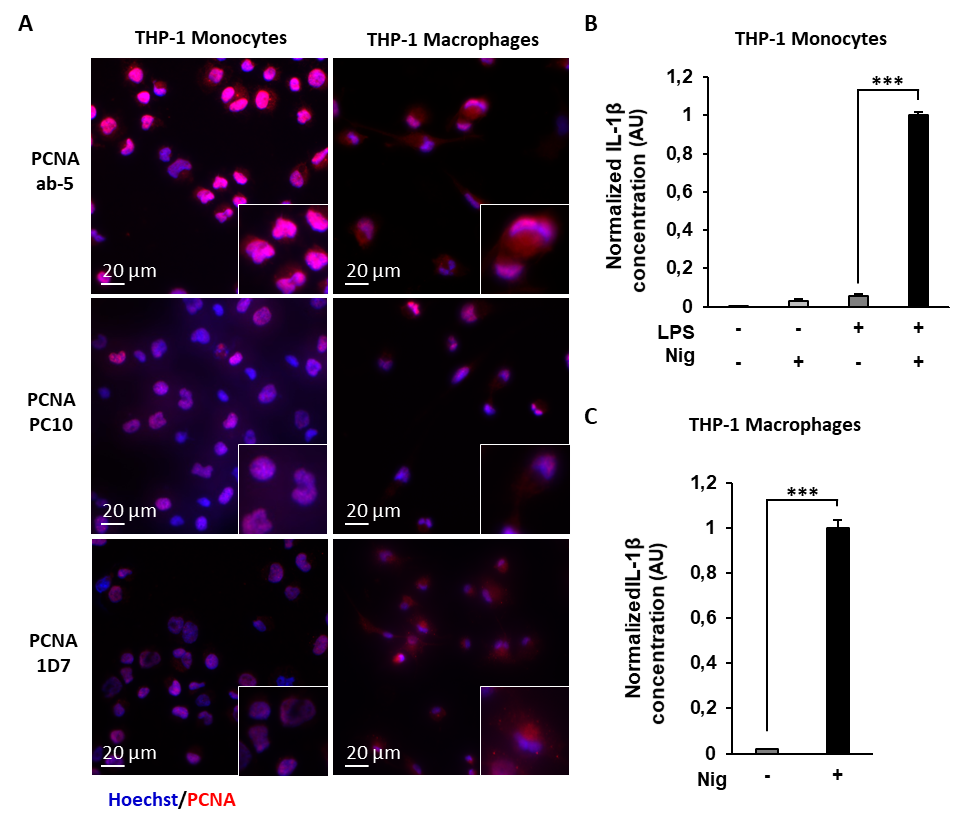

**Supplementary Figure S1**

PCNA expression in THP-1 monocytes and PMA-differentiated THP-1 macrophages. **(A)** Localization of PCNA using PCNA-specific immunofluorescence microscopy analysis performed on monocytic THP-1 cell line or THP-1 cells differentiated into macrophages by PMA (300 ng/ml for 3 h followed by 3 days of resting). Three different anti-PCNA antibodies (ab-5, PC10 and 1D7) were used. Red, PCNA; blue, Hoechst. Representative micrographs are shown. Surrounded cells are magnified 1.8 times. **(B, C)** ELISA quantification of IL-1β secreted by **(B)** THP-1 monocytes cultured for 4 h with (+) or without (−) 200 ng/ml LPS in absence (−) or presence (+) of 20 µM nigericin (Nig) during the last hour of incubation (6 independent experiments performed in duplicate (n=12) or triplicate for LPS+Nig+ (n=14)) or **(C)** PMA-differentiated THP-1 macrophages cultured for 1 h in absence (−) or presence (+) of 20 µM nigericin (Nig) (16 independent experiments performed in duplicate (n=13 ) or triplicate (n=2) or quadruplicate (n=1); total n=36). Results are expressed as mean ± sem of normalized data from independent experiments (AU, arbitrary units). For each independent experiment, results were normalized to the mean of the positive control (LPS+ Nig+ for THP-1 monocytes (ranging from 994,8 to 6551,2 pg/ml) (**Supplementary Figure S2A**) and Nig+ for THP-1 macrophages (ranging from 312,1 to 12177,7 pg/ml) (**Supplementary Figure S2B**)). Non-parametric Mann-Whitney tests were performed, ***p < 0.001.
