## Supplemental Figure 2 for "The cytoplasmic scaffolding protein PCNA regulates NLRP3 inflammasome activation in macrophages"

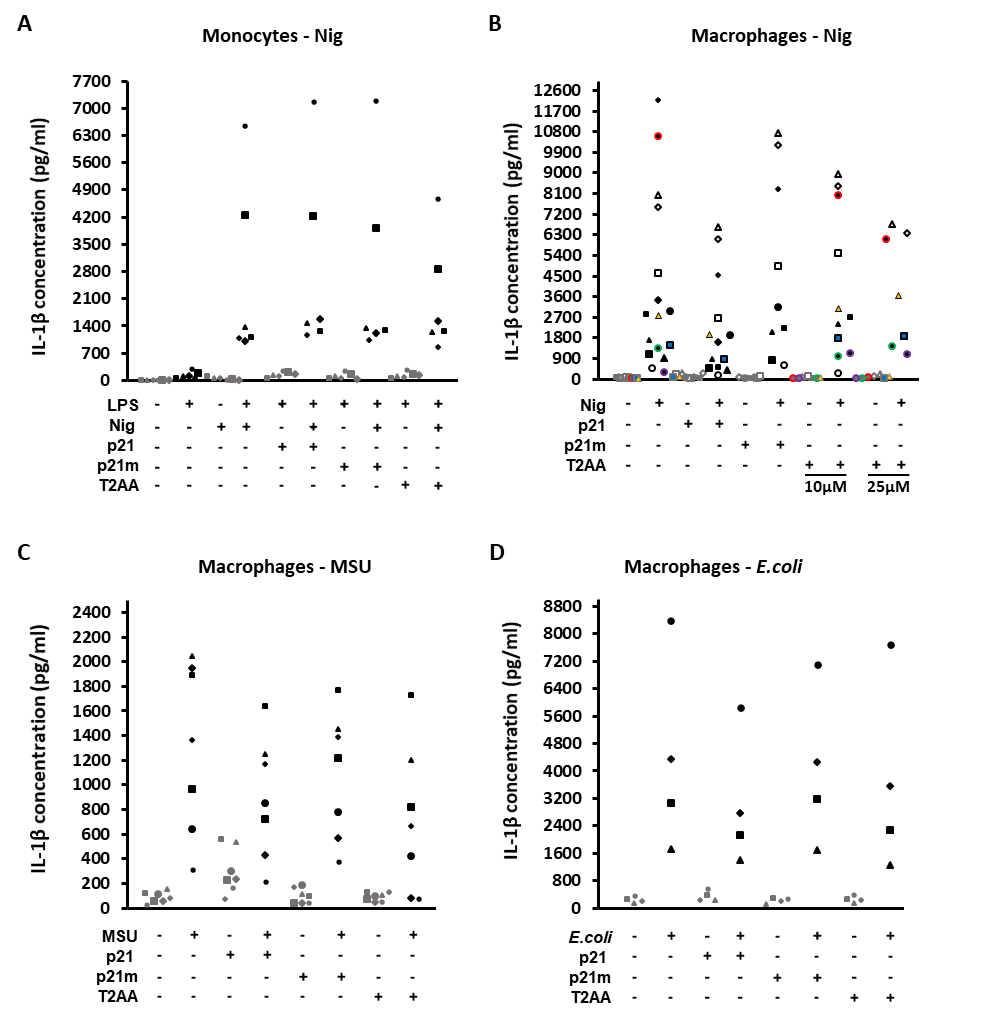

**Supplementary Figure S2**

Preventing the interaction of PCNA with its partners inhibits NLRP3 canonical and non-canonical inflammasome activation in PMA-differentiated THP-1 macrophages. ELISA quantification of IL-1β secreted by **(A)** THP-1 monocytes cultured for 4 h with (+) or without (−) 200 ng/ml LPS pretreated (+) or not (-) with 10 µM p21 peptide (p21) or mutant p21 peptide (p21m) or 25 µM T2AA (6 independent experiments); **(B)** PMA-differentiated THP-1 macrophages pretreated (+) or not (-) (16 independent experiments) with 10 µM p21 (13 independent experiments) or p21m (9 independent experiments), 10 µM T2AA (11 independent experiments) or 25 µM T2AA (7 independent experiments) during 1 h, followed by incubation in absence (−) or presence (+) of 20 µM nigericin (Nig) during 1 h; **(C)** PMA-differentiated THP-1 macrophages pretreated 1h in absence (−) or presence (+) of 10 µM p21, 10 µM p21m or 25 µM T2AA, then cultured for 3 h with (+) or without (−) 200 µg/ml monosodium urate (MSU) crystals (7 independent experiments); **(D)** PMA-differentiated THP-1 macrophages cultured for 2,5 h with (+) or without (−) *E. coli* bacteria at a multiplicity of infection of 20, followed by a 16 h incubation period with penicillin/streptomycin in absence (−) or presence (+) of 5 µM p21, 5 µM p21m or 10 µM T2AA (4 independent experiments). Each point represents the mean IL-1β concentration obtained from one experiment. Each symbol represents an independent experiment.
