## Supplemental Figure 3 for "The cytoplasmic scaffolding protein PCNA regulates NLRP3 inflammasome activation in macrophages"

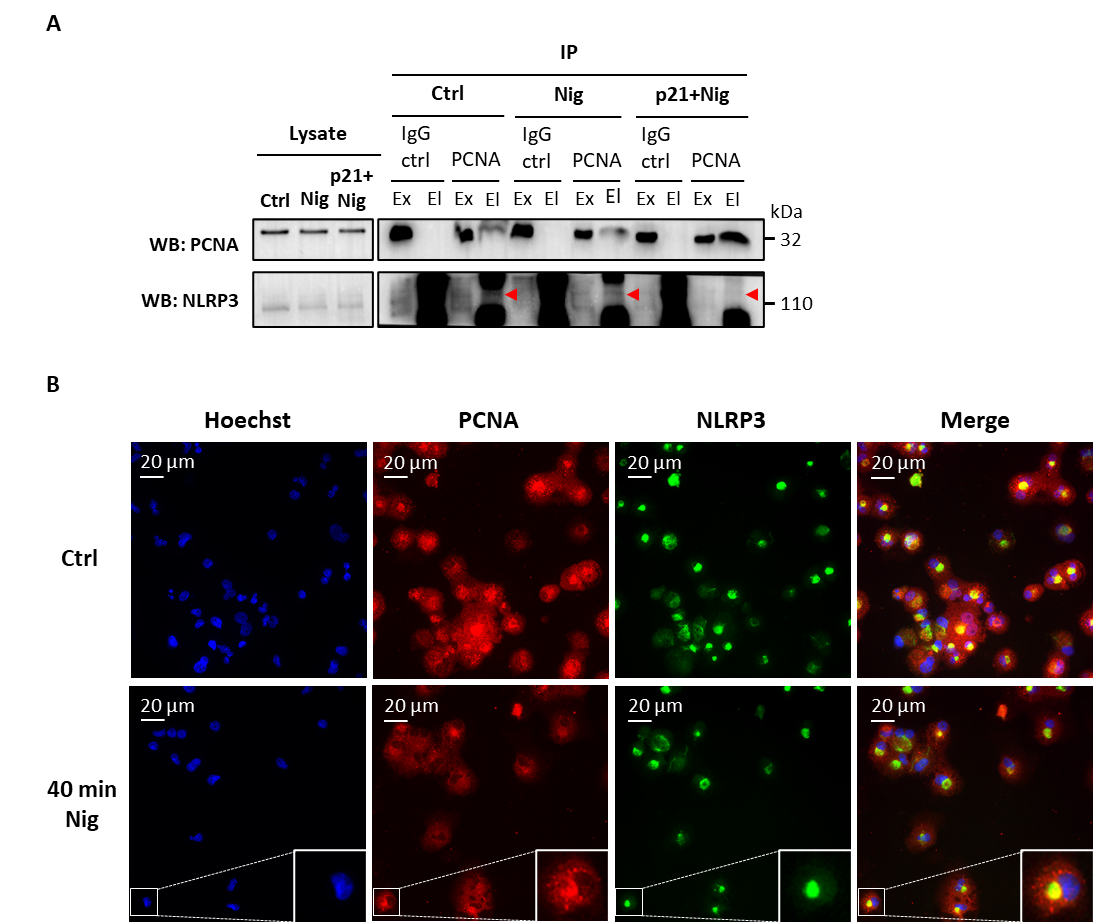

**Supplementary Figure S3**

**(A)** Co-immunoprecipitation (co-IP) experiment were performed on cytosol from PMA-differentiated THP-1 macrophages untreated (ctrl), treated 40 min with 20 µM nigericin (Nig) or pretreated 1 h with 10 µM p21 followed by 40 min incubation with 20 µM nigericin (p21+Nig). Co-IP using beads coated with anti-PCNA pAb (IP: PCNA) or with control isotype Ab (IP: IgG ctrl) were performed. NLRP3 and PCNA were detected by western blot analysis (WB) on the exclusion (Ex) and elution (El) fractions obtained by co-IP. **(B)** Localization of PCNA and NLRP3 using specific immunofluorescence microscopy analysis performed on PMA-differentiated THP-1 macrophages untreated (ctrl, upper panel) or treated 40 min with 20 μM nigericin (Nig, lower panel). Blue, Hoechst; red, PCNA; green, NLRP3. Surrounded cells are magnified 2.5 times.
