## Supplemental Figure 4 for "The cytoplasmic scaffolding protein PCNA regulates NLRP3 inflammasome activation in macrophages"

**SUPPLEMENTARY Materials and methods**

**1. Mice**

p21/Waf1 knockout mice (p21^-/-^) were obtained from The Jackson Laboratory (B6;129S2-Cdkn1atm1Tyj/J; Bar Harbor, ME) and housed with their wild-type (WT) littermate. Mice were bred and maintained in pathogen-free facilities at Cochin Institute, and used between 12-15 weeks of age. All animal experiments followed the European Community Guidelines and were approved by Université Paris Cité Ethics Committee for Animal Research (animal facility agreement C-75-14-02).

**2. Cell culture**

L929 cells were cultured in DMEM with GlutaMAX (Gibco) supplemented with 10% FBS, 1 mM Sodium Pyruvate, 1X NEAA medium (Gibco), 100 U/ml penicillin and 100 μg/ml streptomycin mix. After reaching confluence, L929 cells were cultured 10 additionally days at 37°C in a 5% CO_2_ atmosphere. Conditioned L929 medium is obtained after centrifugation at 1500 g for 10 min to remove any detached cells, filtration (0.22 µM), and is stored at -80°C.

**3. In vitro culture of primary mouse macrophages**

Bone marrow–derived macrophages (BMDM) were obtained by flushing femurs and tibia with DMEM with GlutaMAX containing 10% FBS, 10 mM HEPES, 100 U/ml penicillin and 100 μg/ml streptomycin mix. The bone marrow–cell suspension was filtered using a 40 µm cell strainer and centrifuged at 1300 rpm for 5 min. Erythrocytes were lysed with ACK buffer for 5 min at RT. Lysis was stopped by adding culture medium followed by 10 min centrifugation at 1300 rpm. After cell counting and cell density adjusting, cells were cultured in IMDM, 25 mM HEPES with GlutaMAX (Gibco) supplemented with 10% FBS, 100 U/ml penicillin and 100 μg/ml streptomycin mix, and 20% L929 conditioned medium for 7 days at 37°C in a 5% CO_2_ atmosphere. Adherent cells were used as BMDM. CD11b expression at the BMDM cell surface was confirmed by Fluorescence-Activated Cell Sorting (FACS) analysis. Briefly, 3 x 10^5^ cells were stained for 30 min in the dark with FITC-conjugated anti-CD11b monoclonal antibodies (ebioscience^TM^, clone M1/70) or isotype control antibodies (BD Pharmingen™, FITC-conjugated Rat Anti-Mouse IgG1), then rinsed in PBS, centrifuged for 10 min at 1300 rpm at 4°C, and finally fixed in PBS-2% paraformaldehyde. Data acquisition was carried out using BD Accuri™ C6 Plus, analysed by Cflow Plus software and expressed as the percentage of positive cells as compared to the number of positive cells in the isotype control.

Mouse BMDM were primed with 10 ng/ml LPS overnight in DMEM with 10% FBS, then stimulated in 24-well plates at 10^6^ cells/well in medium without FBS. NLRP3 activation was achieved with 5 µM nigericin for 1 h.

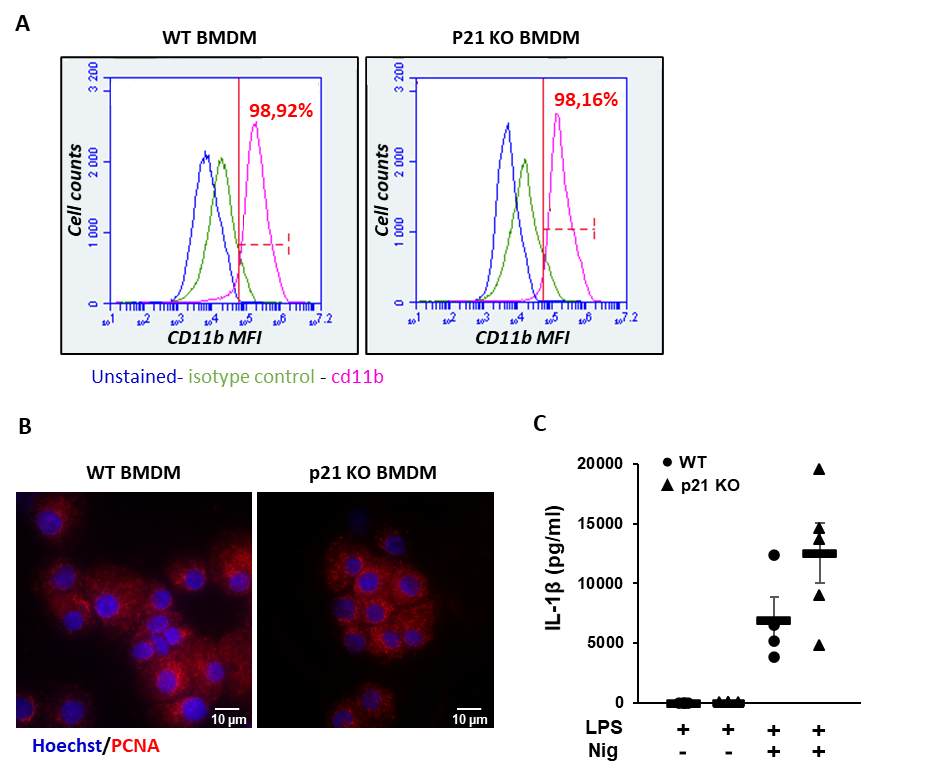

**Supplementary Figure S4**

Increase NLRP3 inflammasome activation in bone marrow derived-macrophages from p21^-/-^mice. **(A)** CD11b expression at the cell surface of bone marrow derived-macrophages (BMDM) from wild type (WT) and p21^-/-^ (p21 KO) mice was detected by FACS analysis. MFI, mean fluorescence intensity. Unstained cells, blue curve; isotype control, green curve; CD11b, pink curve. The Percentage of CD11b-positive cells is expressed as compared to the number of positive cells in the isotype control, and is noted in red. **(B)** Cytoplasmic localization of PCNA in BMDM from WT and p21 KO mice was shown using PCNA-specific immunofluorescence microscopy analysis. Red, PCNA; blue, Hoechst. **(C)** ELISA quantification of IL-1β secreted by WT and p21^-/-^ BMDM cultured for 4 h with (+) 200 ng/ml LPS followed by 2 h incubation in absence (−) or presence (+) of 5 µM nigericin. Each symbol represents one mouse. Bars indicate mean values ± sem.
